# Structural basis for bivalent binding and inhibition of SARS-CoV-2 infection by human potent neutralizing antibodies

**DOI:** 10.1101/2020.10.13.336800

**Authors:** Renhong Yan, Ruoke Wang, Bin Ju, Jinfang Yu, Yuanyuan Zhang, Nan Liu, Jia Wang, Qi Zhang, Peng Chen, Bing Zhou, Yaning Li, Shuyuan Zhang, Long Tian, Xinyue Zhong, Lin Cheng, Xiangyang Ge, Juanjuan Zhao, Hong-Wei Wang, Xinquan Wang, Zheng Zhang, Linqi Zhang, Qiang Zhou

**Author notes:** These authors contributed equally to this work (Renhong Yan, Ruoke Wang, Bin Ju, Jinfang Yu, Yuanyuan Zhang, Nan Liu, Jia Wang, Qi Zhang).

## Abstract

Neutralizing monoclonal antibodies (nAbs) to severe acute respiratory syndrome coronavirus 2 (SARS-CoV-2) represent promising candidates for clinical intervention against coronavirus virus diseases 2019 (COVID-19). We isolated a large number of nAbs from SARS-CoV-2 infected individuals capable of disrupting proper interaction between the receptor binding domain (RBD) of the viral spike (S) protein and the receptor angiotensin converting enzyme 2 (ACE2). In order to understand the mechanism of these nAbs on neutralizing SARS-CoV-2 virus infections, we have performed cryo-EM analysis and here report cryo-EM structures of the ten most potent nAbs in their native full-length IgG or Fab forms bound to the trimeric S protein of SARS-CoV-2. The bivalent binding of the full-length IgG is found to associate with more RBD in the “up” conformation than the monovalent binding of Fab, perhaps contributing to the enhanced neutralizing activity of IgG and triggering more shedding of the S1 subunit from the S protein. Comparison of large number of nAbs identified common and unique structural features associated with their potent neutralizing activities. This work provides structural basis for further understanding the mechanism of nAbs, especially through revealing the bivalent binding and their correlation with more potent neutralization and the shedding of S1 subunit.

## Introduction

The global pandemic of coronavirus disease 2019 (COVID-19) caused by the severe acute respiratory syndrome coronavirus 2 (SARS-CoV-2) is a serious threat to human health(*1, 2*). SARS-CoV-2 is an enveloped, positive-strand RNA virus, belonging to the beta-coronavirus genus that also includes SARS-CoV (*3*) and the Middle Eastern respiratory syndrome coronavirus (MERS-CoV) (*4*) that caused epidemic in 2003 and 2012, respectively. SARS-CoV-2 shares about 80% sequence identity with SARS-CoV, both use angiotensin-converting enzyme 2 (ACE2) as their cellular receptor (*5–9*) that is recognized and bound by the trimeric spike (S) protein (*10, 11*)that distributes on the surface of the virion particles (*12–14*) and is proteolytically cleaved into N-terminal S1 subunit and C-terminal S2 subunit during viral entry into target cells(*15*). S1 contains the N-terminal domain (NTD), the receptor binding domain (RBD), the subdomain 1 and 2 and is responsible for binding to receptor. S2 mediates the fusion of the viral and cellular membrane by undergoing a dramatic conformation change from the prefusion to the postfusion state (*16*) accompanying with the shedding of S1. RBD, which directly binds to ACE2 receptor, is a major target for development of the therapeutic neutralizing monoclonal antibodies (nAbs) against COVID-19. The prefusion structure of S protein exhibits more dynamic conformational changes in S1 region, especially in RBD, which has two distinctive conformations: “up” and ‘‘down”(*10, 11*). Only the “up” conformation of RBD can bind to the ACE2 receptor. Up to now, numerous nAbs against the S protein of SARS-CoV-2 have been reported (*17–23*). The complex structures of these nAbs with S protein were solved, most of which utilized the Fab form of nAbs. It remains largely unknown how nAbs in their native bivalent form bind to and ever induce the conformation changes of the trimeric S protein.

To further explore the interactions between nAbs and S proteins, we solved ten cryo electron microscopy (cryo-EM) structures of the S protein in complex with nAbs, in full-length form and/or in Fab form. Bivalent binding was revealed for the full-length form nAbs, which showed that the full-length form exhibits different binding mode and induces more RBDs to the “up” conformation than the Fab form that is monovalent. It is also shown that the bivalent binding is superior in antiviral efficacy, and correlated with the improved shedding of S1 subunit. Structural comparison of large number of the complexes of nAbs with S protein identified common and unique features associated with the potent neutralizing activities of these nAbs. Our results provide an important structural basis for further understanding the working mechanism of nAbs and are helpful for antiviral drug design and vaccine development.

### Potent nAbs isolated from the COVID-19 convalescent patients

To understand the molecular features of the interactions of neutralizing nAbs with S protein, we characterized ten nAbs derived from various COVID-19 convalescents about the binding and neutralizing activities, and the capacity of competing with ACE2 for RBD binding. The binding affinity to RBD of SARS-CoV-2 of these nAbs measured by surface plasmon resonance varied from 0.75 to 90.09 nM (Fig. S1A and Table 1), whereas the half maximal inhibitory concentration (IC_50_) of these nAbs ranged from 0.0014 to 0.9231 μg/ml in the pseudo virus based assay or from 0.0043 to 0.8932 μg/ml in live SARS-CoV-2 virus based assay, respectively (Ref, Table 1 and Fig. S1). All of these nAbs exhibited strong competition with ACE2 to bind RBD, indicating their neutralization mechanism (Table 1 and Fig. S1). The variable regions of the heavy chain of these nAbs belong to diverse gene families, paired with different families of light chains. The CDR3 length of the heavy chains and the light chains ranged from 9 to 22 amino acids and from 9 to 11 amino acids, respectively (Table S1). The somatic hypermutation (SHM) of these nAbs were generally low and five of them reached 0% for heavy chain or light chain.

**Table 1.** Binding capacity, neutralizing activity, and gene family analysis of COVID-19 donor-derived neutralizing nAbs. * Published in the reference (Zhang, et al. Public neutralizing antibodies elicited by SARS-CoV-2 infection. submitted). The program IMGT/V-QUEST was applied to analyze gene germline, complementarity determining region (CDR) 3 length, and somatic hypermutation (SHM). The CDR3 length was calculated from amino acids sequences. The SHM frequency was calculated from the mutated nucleotides. Antibody binding to RBD was presented either by Kd or by competing with ACE2 where “+++” indicates >80% competition. IC_50_ represents the half-maximal whereas IC_80_ the 80% inhibitory concentrations in the pseudovirus and live SARS-CoV-2 neutralization assay.

| mAbs | Binding to RBD |  | Pseudovirus (µg/ml) |  | Live virus (µg/ml) |  | Shedding at 120min |
| --- | --- | --- | --- | --- | --- | --- | --- |
|  | Kd (nM) | competing w/ ACE2 | IC<sub>50</sub> * | IC<sub>80</sub> * | IC<sub>50</sub> * | IC<sub>80</sub> * |  |
| P2B-1A10* | 50.77 | +++ | 0.0974 | 0.7446 | 0.0639 | 0.3053 | 81.80% |
| P5A-3A1 | 90.09 | +++ | 0.9231 | 4.2357 | 0.6713 | 26.2193 | 77.20% |
| P5A-1B8* | 1.09 | +++ | 0.0115 | 0.0501 | 0.0168 | 0.0857 | 79.60% |
| P5A-2G9* | 6.98 | +++ | 0.0158 | 0.1466 | 0.0113 | 0.1187 | 84.90% |
| P5A-1B6 | 1.01 | +++ | 0.2528 | 1.3719 | 0.8932 | 5.9133 | 76.00% |
| P2B-1A1 | 26.97 | +++ | 0.6900 | 2.4100 | 0.2218 | 2.1498 | -9.70% |
| P5A-2G7* | 3.55 | +++ | 0.0044 | 0.0287 | 0.1814 | 0.8355 | 57.20% |
| P5A-1B9* | 0.75 | +++ | 0.0014 | 0.0053 | 0.0043 | 0.0441 | 43.80% |
| P5A-2F11 | 5.33 | +++ | 0.6300 | 1.9400 | 0.4942 | 6.9416 | 56.20% |
| P5A-3C12* | 1.03 | +++ | 0.0996 | 0.4679 | 0.2636 | 2.6783 | 54.50% |

\* Published in Zhang, et al (submitted)

Inducing shedding of the surface proteins of viruses is an important feature for nAbs. So, we decided to examine whether the nAbs in this work can induce the shedding of the S1 subunit of the S protein of SARS-CoV-2 or not by transfecting plasmids encoding SARS-CoV-2 S protein into HEK293T cells. The results showed that some nAbs, such as P5A-2G9, could notably induce shedding of the S protein, whereas some nAbs, such as P5A-1B9 and P5A-3C12, had only a weak shedding ability and P2B-1A1 exhibited almost no shedding ability, suggesting the diversity of these nAbs in terms of inducing the shedding of S1 subunit.

### The complex structures of the SARS-CoV-2 S trimer bound with nAbs

To get deeper understanding of the working mechanism, we determined the complex structures of the full-length IgG form of these nAbs with the S protein of SARS-CoV-2 using single particle cryo-EM at overall resolution from 2.8 angstroms to 3.6 angstroms (Fig. 1, Fig. S2-6 and Table S2). The nAbs were incubated with the S protein at excessive molar ratio and the unbound nAbs were removed by gel filtration (Fig. S2). Focused refinement at the interface between nAbs and RBD of the S protein notably improved the map quality, allowing detailed analysis. Additionally, we also prepared the cryo-EM samples of the S protein incubated with Fab region of P5A-3A1, P5A-3C12 in a short period of time. For the Fab-bound complexes, only the structures with lower resolution at the interface of Fab and RBD were obtained (Fig. 1, Fig. S7-8). For clarity, we use IgG and Fab to indicate the bivalent full-length form and the monovalent Fab form of nAbs, respectively. If there is no such specific label, it means the bivalent full-length form of nAb.

**Fig. 1.**
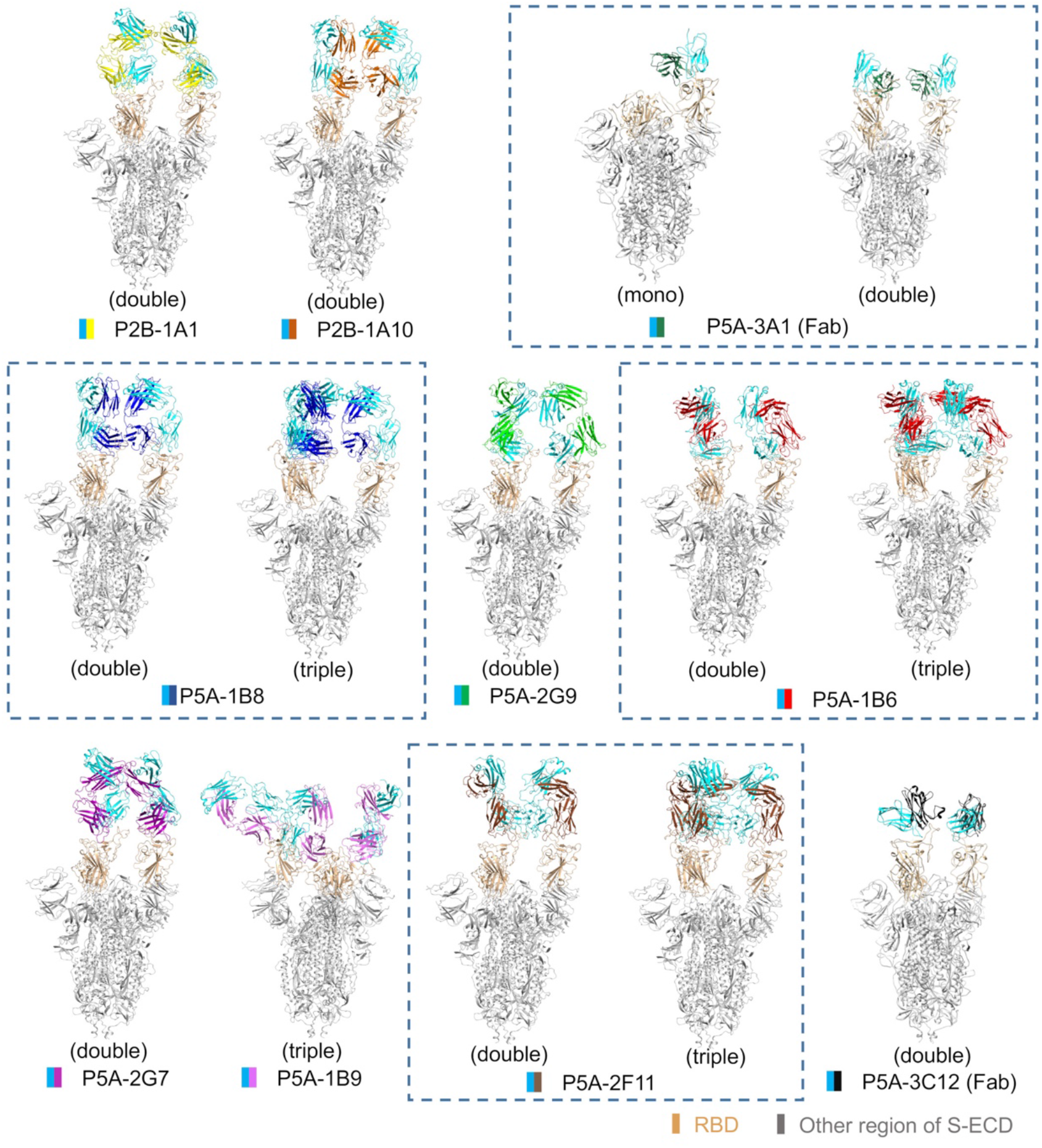
All solved structures of nAbs in complex with S protein. The domain-colored models of the all complex are shown here. The structures containing different number of the same nAb are boxed with blue dash line. The structures are labelled according to the number of RBD bound with nAb as mono (1 RBD), double (2 RBDs) or triple (3 RBDs) binding, respectively.

The structures of these S-IgG complexes can be classified into three different binding patterns (Fig. 1). In pattern 1 that includes P2B-1A10, P2B-1A1, P5A-2G7 and P5A-2G9, two “up” RBDs are bound with nAb. In pattern 2 that contains P5A-2F11, P5A-1B8 and P5A-1B6, two or three RBDs are in “up” conformation and bound with nAb. P5A-1B9, the most potent nAb in this work, itself constitutes the pattern 3 that contains one “up” RBD and two “down” RBD. All of three RBDs in P5A-1B9 are bound with nAb and share the same binding interface (Fig. S9).

### The bivalent binding of nAbs

It’s reported that the bivalent binding of antibodies can neutralize the virus more efficiently than Fabs in some viruses such as rhinovirus and Dengue virus (*24, 25*). To examine the structural difference of the complexes of the S protein with IgG and with Fab, we further solved the complex structures of the S protein with Fab region of P5A-1B8 or P5A-2G7 (Fig. 2, Fig. S8). In the S/P5A-1B8(IgG) complex, two or three RBDs are in “up” conformation and bound with nAb (Fig. 1), whereas only two RBDs are in “up” conformation and bound with nAb in the S/P5A-1B8(Fab) complex. Next we choose the S/P5A-1B8(IgG) complex containing two “up” RBDs to compare with the S/P5A-1B8(Fab) complex. The structural comparison showed that the two cryo-EM densities corresponding to the nAb in the S/P5A-1B8(IgG) complex are closer to each other than that in the S/P5A-1B8(Fab) complex (Fig. 2A). When one of the Fab regions of the S/P5A-1B8(IgG) complex is superimposed with that of the S/P5A-1B8(Fab) complex, the other Fab exhibits different orientation (Fig. 2A). For P5A-2G7, only one Fab binds to one “up” RBD in the S/P5A-2G7(Fab) complex, whereas two “up” RBDs are bound with nAb in the S/P5A-2G7(IgG) complex, indicating a different binding pattern between the IgG form and the Fab form (Fig. 2B).

**Fig. 2.**
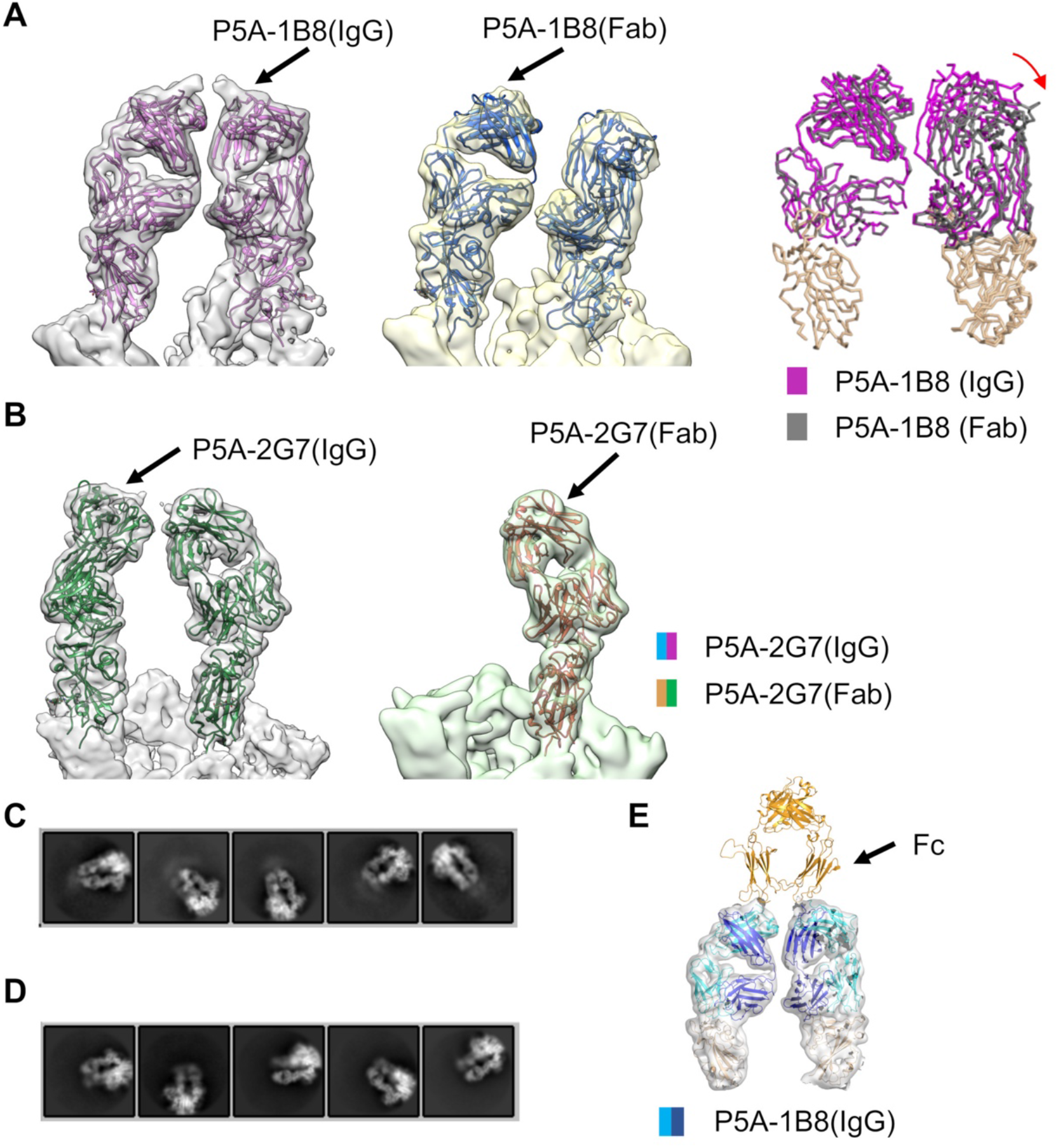
Bivalent binding analysis of nAbs. **A**, Structural comparison between the S/P5A-1B8(IgG) complex and the S/P5A-1B8(Fab) complex. The cryo-EM maps docked with atomic models are shown in left and middle panels. Right panel: The atomic models were superimposed according to one arm to show the difference in the other arm. **B,** Structural comparison between S-P5A-2G7 and S-Fab-2G7. Right panel: The interfaces alignment. **C** and **D** are 2D classification of the S/P5A-1B8(IgG) complex and the S/P5A-1B8(Fab) complex re-centered at antibody, respectively. **E,** The cryo-EM map of the S/P5A-1B8(IgG) complex can be docked with the Fc region of a full-length antibody model. The Fc, heavy chain, and light chain of the antibody are colored in gold, blue and cyan, respectively.

We also re-centered the particles of the S/P5A-1B8(IgG) complex and the S/P5A-1B8(Fab) complex at the Fab region and calculated the two-dimensional (2D) classification. The 2D class averages showed extra density near nAb of the S/P5A-1B8(IgG) complex, which might correspond to the Fc region of the P5A-1B8 (Fig. 2C) and was absent in the 2D class averages of the S/P5A-1B8(Fab) complex (Fig. 2D). Additionally, the Fab regions of an intact antibody molecule can be docked into the cryo-EM map of the S/P5A-1B8(IgG) complex (Fig. 2E). Taken together, these results support the bivalent binding exists for both P5A-1B8 and P5A-2G7.

### IgG-form nAbs shows advantage over Fab-form ones in neutralizing potency and shedding of S1 subunit

To investigate the working mechanism more deeply, we compared the neutralizing potency and the ability to induce the shedding of S1 between the IgG form and the Fab form of the nAbs. As shown in Fig. 3A, the IgG form nAbs exhibited higher neutralizing potency than the Fab-form of the same nAbs. The neutralizing IC_50_ values of the IgG-form were decreased for 700-fold compared to the Fab form for P5A-1B8 and P5A-1B9, whereas P5A-2G7 exhibited even greater decreasing of more than 3,000 folds. To investigate the ability of inducing the shedding of S1 subunit for these nAbs in IgG- or Fab-forms, we incubated S protein-surface-expressed cells with IgG or Fab at saturated concentration and measured their binding over time by flow cytometry. As shown in Fig. 3B, the IgG of P5A-1B8 triggered S1 shedding most potently, which induced the shedding of S1 subunit for about 80% of the S protein after incubating with cells for 120 minutes. However, the Fab-form of P5A-1B8 nearly completely lost the ability of inducing the shedding of S1. P5A-1B9 and P5A-2G7 showed mild shedding ability. As a control and consistent with the expectation, S1 of the GSAS-containing mutant cannot be shed by either IgG or Fab (Fig. 3C). The rest nAbs tested also exhibit similar properties (Fig. S10).

**Fig. 3.**
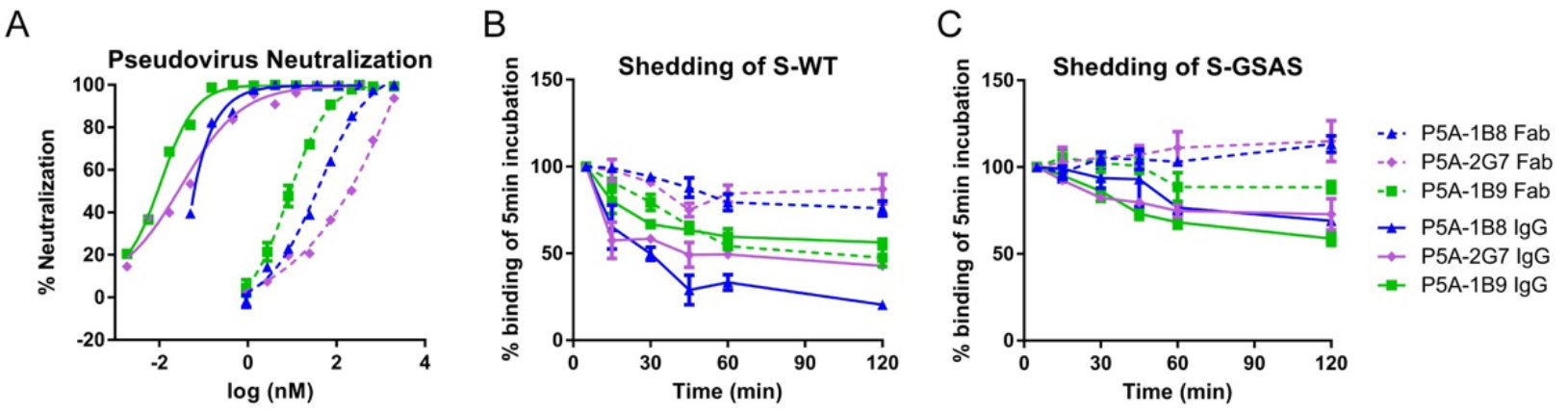
Neutralizing activity and shedding of S1 by IgG and Fab forms of nAbs. **A**, Neutralizing activity against SARS-CoV-2 pseudovirus by P5A-1B8, P5A-1B9, and P5A-2G7 in IgG forms (solid line) and Fab forms (dotted line). **B** is the shedding of S1 over time measured using flow cytometry at 37°C with 293T cell-surface expressed wild type SARS-Cov-2 S protein. **C** is same to B, with a mutant S protein containing GSAS substitution at S1/S2 cleavage motif. Data shown were from at least two independent experiments.

### Comparisons of antibody binding epitopes

A summary of the above cryo-EM structure determinations enabled us to compare and classify the epitopes of these antibodies (Fig. 4 and Fig. S11). In the final models, the Fab was successfully built for antibodies P2B-1A10, P5A-1B8, P5A-2G9, P5A-1B6, P2B-1A1, P5A-2G7, P5A-1B9 and P5A-2F11, and scFv was built for antibodies P5A-3A1 and P5A-3C12 due to the weak densities in the CH1 domains. These ten neutralizing antibodies all target the receptor binding motif (RBM) of RBD and could be classified into three groups, considering the epitopes and approaching angles to the RBD. The first group, including P2B-1A10, P5A-3A1, P5A-1B8, P5A-2G9, P5A-1B6, P2B-1A1 and P5A-2G7, has the largest overlap between the epitope and RBM. Their epitope residues are distributed across the RBM, mainly in the cradle region. Among the 17 residues of RBD involved in receptor binding, more than half (8 to 15) were recognized by the antibodies in the group 1. The antibodies in group 1 also have similar contacting angles with the RBD ranging from 39 to 52 degrees, and can be further divided into three subgroups. The subgroup 1 consists of three antibodies P2B-1A10, P5A-3A1 and P5A-1B8, which use the same heavy chain IGHV3-53 V gene and exhibit very similar positional arrangement. They are all antibodies whose heavy chain plays a leading role. Among the 17 residues of RBD involved in receptor binding, 8 to 9 were recognized by the heavy chains in the subgroup1 while the number is 2 to 5 by light chains. The shedding activity is higher than 77.2%. When RBD of the structures in first subgroup was aligned, the distances of VH domains or VL domains are ranging from 1.8 to 2.3 angstroms or from 2.4 to 3.3 angstroms, respectively. The usage of the IGHV3-53/ IGHV3-66 V genes have also been reported for other antibodies such as B38 and CB6 (*23, 26*).

**Fig. 4.**
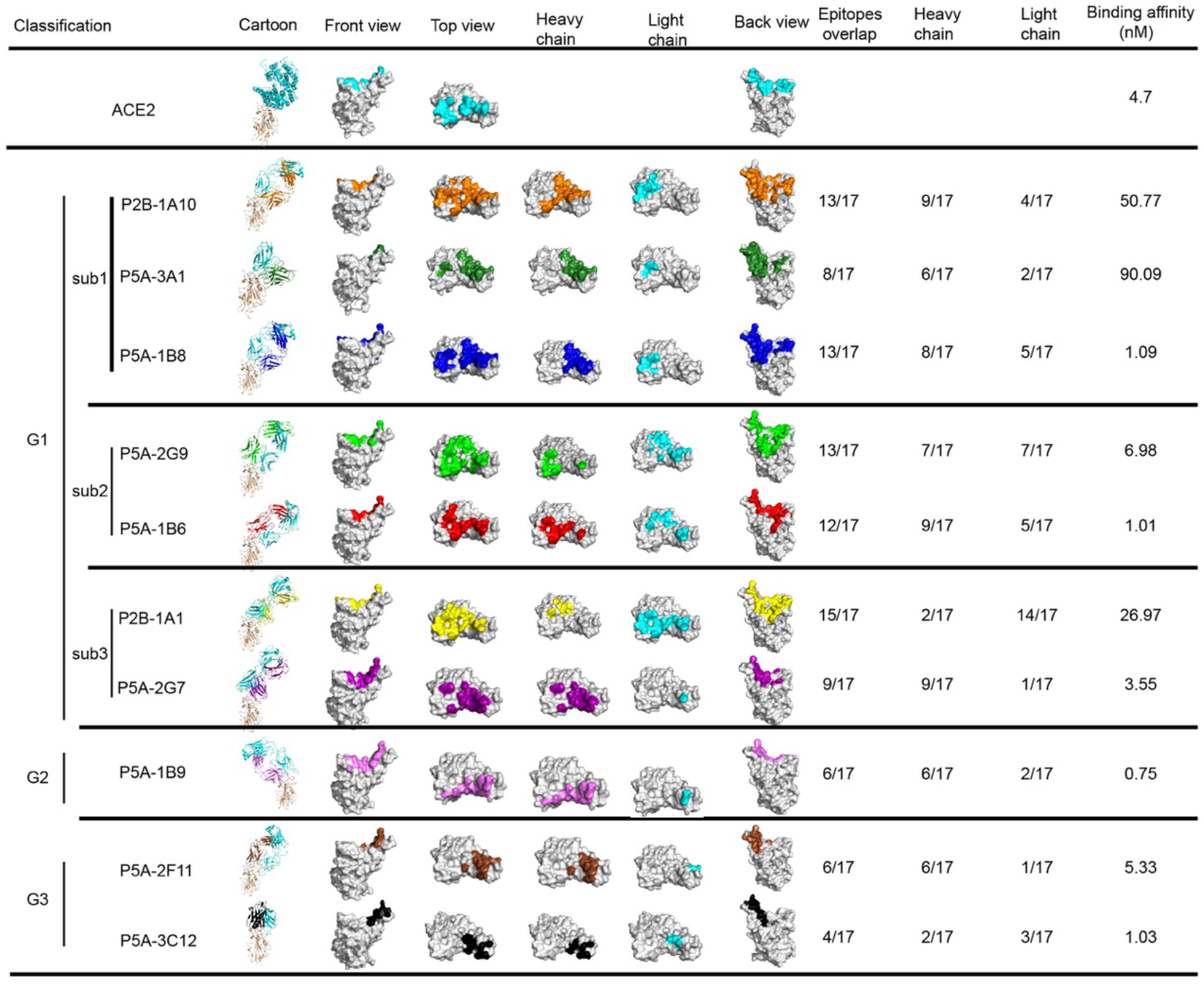
The epitopes of 10 anti SARS-CoV-2 RBD nAbs. The 10 antibodies could be classified into three groups. The first group antibodies can be further divided into three subgroups. The complexes of RBD with ACE2 or nAbs are shown as cartoon with RBD colored in wheat, the light chains of nAbs colored in cyan and the heavy chains of nAbs colored in different colors. RBD is also shown as grey surface in top, front and back views, with the RBM colored in cyan and the epitopes of different nAbs shown in respective colors. For the top views, the epitopes corresponding to heavy chains and lights chains are shown separately in respective colors for heavy chains or in cyan for light chains.

The subgroup 2 contains P5A-2G9 and P5A-1B6 and uses heavy chain gene from IGHV 3 family, but not IGHV3-53 gene. The subgroup 2 has a different VH and VL positional arrangement from the subgroup 1. The subgroup 3 contains P2B-1A1 and P5A-2G7, which use heavy chain gene from IGHV4 family and have a rotation around the longitudinal axis of the Fab compared with the antibodies in subgroup 1 and 2. The heavy chain of P5A-2G7 plays the leading role like subgroup1 and subgroup2 antibodies while the light chain of P2B-1A1 plays the leading role.

The P5A-1B9 alone forms the group 2 among the eight neutralizing antibodies we have structurally characterized. The inter-molecular angle between P5A-1B9 and the RBD is 52 degrees anti-clockwise while the ACE2 degree is clockwise. Upon binding, P5A-1B9 approaches the RBD from a direction different from those of the first group antibodies. The epitope for P5A-1B9 has 6 residues overlapping with RBM. The heavy chain contributes 6 residues and the light chain contribute 2 residues. The shedding ability is 43.8%. The group 3 consists of antibodies P5A-2F11 and P5A-3C12, which use heavy chain genes from IGHV1 and IGHV2 family, respectively. The epitopes for antibodies in group 3 are mainly located in the remote loops and less overlapped with RBM, with only 6 or 4 overlapping residues for P5A-2F11 or P5A-3C12, respectively. The heavy chain of P5A-2F11 plays the leading role while the light chain of P5A-3C12 plays the leading role. The shedding ability is higher than 54.5%.

## Discussion

In this work, we solved ten complex structures of the S protein with nAbs in full-length IgG form and/or in Fab form. Structural analysis revealed the bivalent binding mode of nAbs against SARS-CoV-2. Our biochemical and cell-based experimental results showed that the full-length IgG nAbs have greater binding ability, neutralization ability, and induce more S1 shedding than Fab. Similar results were also obtained in other study(*27*). Moreover, there are more RBDs in “up” conformation in the structures of the S protein in complex with full-length IgG than that in complex with Fab. The difference should be attributed to the bivalent binding of IgG molecules that contain two Fab molecules. The full-length IgG nAbs bind to SARC-CoV-2 S proteins in bivalent mode with higher probability than the Fabs thus may induce stronger neutralization. Furthermore, the cooperation of the RBD conformational change induced by the bivalent binding would enhance the induced-conformational change of the S proteins on a virus surface. More importantly, the presence of Fc region in the antibody-virus complex can enhance the immunity response by the Fc receptors of T cells. Up to now, numerous complex structures of the S protein with nAb have been reported, most of which used nAb in Fab-form, especially those solved with X-ray crystallography. Our results suggest that the full-length IgG-form, which is more biologically relevant than Fab, should be used for the structural determination as much as possible to understand the structural and functional features of nAbs.

Among the ten nAbs reported in this research, the subgroup 1 and 2 of group 1 have a higher shedding ability (above 76 %), while the subgroup 3 of group 1, group 2 and group 3 show no or a weaker shedding ability (43.8% - 57.2%). The group 1 has the largest overlap between the epitope and RBM, while the members in subgroup 3 have a rotation around the longitudinal axis of the Fab. The S1 shedding ability of the nAbs may be facilitated by the large overlap with RBM and require special angle to bind with RBD.

The SARS-CoV and SARS-CoV-2 cross-reactive antibodies that show neutralizing activity mainly target to RBD, interfere with ACE2 binding and stimulate S1 dissociation (*28*), suggesting the ability to induce the shedding of S1 is correlated with the potency of the neutralizing nAbs. The shedding of S1 might be facilitated by the RBD in up conformation that is stabilized by the binding of nAb or receptor. It is not clear whether the shedding of all of three S1 subunits is required for the transition from the prefusion state to the postfusion state to catalyze the fusion of viral and cellular membrane or not. Our previous work indicates the receptor ACE2 exists as a dimer. Each protomer of ACE2 dimer can be bound with one RBD from a S protein trimer and induces the bound RBD to the “up” conformation. It seems unlikely that one ACE2 dimer binds to a trimeric S protein simultaneously because of the steric clash. To induce more than one RBDs in a S protein to up conformation, multiple copies of ACE2 are required.

Up to now, there are 17 structures involving anti-SARS-CoV-2 nAbs in the Protein Data Bank (PDB) database. The group 1 nAbs, which have the largest overlap between the epitope and RBM, not only are most popular in this work, but have many similar nAbs in the PDB database, including B38 (PDB code: 7BZ5)(*26*), CB6 (PDB code: 7C01)(*23*), C105 (PDB code: 6XCN)(*29*), CV30 (PDB code: 6XE1)(*30*), CC12.1 (PDB code: 6XC2)(*31*), CC12.3 (PDB code: 6XC4)(*31*), COVA2-04 (PDB code: 7JMO)(*32*) and COVA2-39 (PDB code: 7JMP)(*32*). Interestingly, the P2B-1A10, P5A-3A1, P5A-1B8, B38, C105, CV30, CC12.1, CC12.3, COVA2-04 and COVA2-39 are all belong to the IGHV3-53 gene family. The group 2 nAb P5A-1B9 has the highest inhibitory activities against the cell infection of both pseudo and live SARS-CoV-2. Structure of the complex of P5A-1B9 with S protein shows that it can bind to RBD in both up and down conformation, similar to the nAb BD-368-2 reported by other group (*33*), suggesting a common mechanism behind these nAbs of very high potency. The epitope of P5A-1B9 is only 6-residue overlapped with RBM, and is similar to P2B-2F6 (PDB code: 7BWJ)(*17*), Fab2-4 (PDB code: 6XEY)(*34*) and BD23 (PDB code: 7BYR)(*35*), but they have different gene family. P2B-2F6 and Fab2-4 have different approaching direction between the nAbs and RBD to the group 1 nAbs. The group 3 nAbs, P5A-2F11 and P5A-3C12, only bind to the remote loop of RBM and there is no published nAb of the similar binding mode.

## Acknowledgments

We thank the cryo EM facility and the supercomputer center of Westlake University and the cryo-EM facility and the bio-computing platform at Tsinghua University Branch of China National Center for Protein Sciences (Beijing) for providing cryo-EM and computation supports. This work was funded by the National Natural Science Foundation of China (projects 31971123, 81920108015, 31930059, 81530065, 91442127, 82002140), the Key R&D Program of Zhejiang Province (2020C04001), the SARS-CoV-2 emergency project of the Science and Technology Department of Zhejiang Province (2020C03129), the Beijing Advanced Innovation Center for Structural Biology and the National Key Plan for Scientific Research and Development of China (2020YFC0848800, 2020YFC08442), the National Key Plan for Scientific Research and Development of China (2020YFC0848800 and 2020YFC0844200), the Science and Technology Innovation Committee of Shenzhen Municipality (202002073000002, 2020A1111350032, JCYJ20190809115617365, and 2020B1111340074), the Natural Science Foundation of Guangdong Province of China (2019A1515011197), Beijing Municipal Science and Technology Commission (Z201100005420019), the Leading Innovative and Entrepreneur Team Introduction Program of Hangzhou, and Special Research Program of Novel Coronavirus Pneumonia of Westlake University and Tencent Foundation, Tsinghua University Initiative Scientific Research Program (20201080053). We would like to express our sincere gratitude towards the generous supports from Tencent Foundation, Shuidi Foundation, and TH Capital.

## Author contributions

Q. Zhou, L.Z., Z.Z., X.W., and H.W. conceived the project. R.Y., R.W., J.Y., Y.Z., N.L., B.J., J.W., Q. Zhang, P.C., B.Z., S.Z., L.T., X.Z., L.C. X.G. and J.Z. did the experiments. All authors contributed to data analysis. R.Y., R.W., J.Y., H.W., L.Z. and Q. Zhou wrote the manuscript.

## Data availability

Atomic coordinates and cryo EM density maps of the nAbs in complex with S protein have been deposited to the Protein Data Bank (http://www.rcsb.org) and the Electron Microscopy Data Bank (https://www.ebi.ac.uk/pdbe/emdb/), respectively. Please refer to Table S2 for the PDB and EMDB codes. Correspondence and requests for materials should be addressed to.

## References

1. F. Wu et al., A new coronavirus associated with human respiratory disease in China. Nature 579, 265–269 (2020).

2. N. Zhu et al., A Novel Coronavirus from Patients with Pneumonia in China, 2019. N Engl J Med 382, 727–733 (2020).

3. T. G. Ksiazek et al., A novel coronavirus associated with severe acute respiratory syndrome. N Engl J Med 348, 1953–1966 (2003).

4. A. M. Zaki, S. van Boheemen, T. M. Bestebroer, A. D. Osterhaus, R. A. Fouchier, Isolation of a novel coronavirus from a man with pneumonia in Saudi Arabia. N Engl J Med 367, 1814–1820 (2012).

5. J. Lan et al., Structure of the SARS-CoV-2 spike receptor-binding domain bound to the ACE2 receptor. Nature, (2020).

6. J. Shang et al., Structural basis of receptor recognition by SARS-CoV-2. Nature, (2020).

7. Q. Wang et al., Structural and Functional Basis of SARS-CoV-2 Entry by Using Human ACE2. Cell, (2020).

8. R. Yan et al., Structural basis for the recognition of the SARS-CoV-2 by full-length human ACE2. Science, (2020).

9. W. Li et al., Angiotensin-converting enzyme 2 is a functional receptor for the SARS coronavirus. Nature 426, 450–454 (2003).

10. A. C. Walls et al., Structure, Function, and Antigenicity of the SARS-CoV-2 Spike Glycoprotein. Cell, (2020).

11. D. Wrapp et al., Cryo-EM structure of the 2019-nCoV spike in the prefusion conformation. Science 367, 1260–1263 (2020).

12. Z. Ke et al., Structures and distributions of SARS-CoV-2 spike proteins on intact virions. Nature, (2020).

13. H. Yao et al., Molecular architecture of the SARS-CoV-2 virus. bioRxiv, (2020).

14. B. Turonova et al., In situ structural analysis of SARS-CoV-2 spike reveals flexibility mediated by three hinges. Science, (2020).

15. M. Hoffmann et al., SARS-CoV-2 Cell Entry Depends on ACE2 and TMPRSS2 and Is Blocked by a Clinically Proven Protease Inhibitor. Cell, (2020).

16. Y. Cai et al., Distinct conformational states of SARS-CoV-2 spike protein. Science, (2020).

17. B. Ju et al., Human neutralizing antibodies elicited by SARS-CoV-2 infection. Nature 584, 115–119 (2020).

18. X. Chi et al., A neutralizing human antibody binds to the N-terminal domain of the Spike protein of SARS-CoV-2. Science 369, 650–655 (2020).

19. P. J. M. Brouwer et al., Potent neutralizing antibodies from COVID-19 patients define multiple targets of vulnerability. Science 369, 643–650 (2020).

20. D. Wrapp et al., Structural Basis for Potent Neutralization of Betacoronaviruses by Single-Domain Camelid Antibodies. Cell 181, 1004–1015.e1015 (2020).

21. M. Yuan et al., Structural basis of a shared antibody response to SARS-CoV-2. Science, (2020).

22. J. Hansen et al., Studies in humanized mice and convalescent humans yield a SARS-CoV - 2 antibody cocktail. Science, (2020).

23. R. Shi et al., A human neutralizing antibody targets the receptor-binding site of SARS-CoV-2. Nature 584, 120–124 (2020).

24. E. A. Hewat, D. Blaas, Structure of a neutralizing antibody bound bivalently to human rhinovirus 2. EMBO J 15, 1515–1523 (1996).

25. M. A. Edeling et al., Potent dengue virus neutralization by a therapeutic antibody with low monovalent affinity requires bivalent engagement. PLoS Pathog 10, e1004072 (2014).

26. Y. Wu et al., A noncompeting pair of human neutralizing antibodies block COVID-19 virus binding to its receptor ACE2. Science 368, 1274–1278 (2020).

27. B. Wang et al., Bivalent binding of a fully human IgG to the SARS-CoV-2 spike proteins reveals mechanisms of potent neutralization. bioRxiv, (2020).

28. A. Z. Wee et al., Broad neutralization of SARS-related viruses by human monoclonal antibodies. Science 369, 731–736 (2020).

29. C. O. Barnes et al., Structures of Human Antibodies Bound to SARS-CoV-2 Spike Reveal Common Epitopes and Recurrent Features of Antibodies. Cell 182, 828–842 e816 (2020).

30. N. K. Hurlburt et al., Structural basis for potent neutralization of SARS-CoV-2 and role of antibody affinity maturation. bioRxiv, (2020).

31. M. Yuan et al., Structural basis of a public antibody response to SARS-CoV-2. bioRxiv, (2020).

32. N. C. Wu et al., An alternative binding mode of IGHV3-53 antibodies to the SARS-CoV-2 receptor binding domain. bioRxiv, (2020).

33. S. Du et al., Structures of potent and convergent neutralizing antibodies bound to the SARS-CoV-2 spike unveil a unique epitope responsible for exceptional potency. bioRxiv, (2020).

34. L. Liu et al., Potent neutralizing antibodies against multiple epitopes on SARS-CoV-2 spike. Nature 584, 450–456 (2020).

35. Y. Cao et al., Potent Neutralizing Antibodies against SARS-CoV-2 Identified by High-Throughput Single-Cell Sequencing of Convalescent Patients’ B Cells. Cell 182, 73–84 e16 (2020).

