## Supplemental Information for "Structural basis for bivalent binding and inhibition of SARS-CoV-2 infection by human potent neutralizing antibodies"

### Materials and methods

#### Antibody and Fab fragment production

Antibody production was conducted as previously described. Briefly, genes encoding the heavy and light chains of antibodies were transiently transfected into HEK 293F cells using polyethylenimine (PEI) (Sigma). After 96h, antibodies in the supernatant were collected and captured by Magnetic Protein A beads (Genscript). Bound antibodies were eluted and further purified by gel filtration chromatography using a Superdex 200 High Performance column (GE Healthcare). To produce Fab fragments, antibodies were cleaved using Protease Lys-C (Roche) with an IgG to Lys-C ratio of 4000:1 (w/w) in 10mM EDTA, 100mM Tris-HCl, pH 8.5 at 37 °C for approximately 12h. Fc fragments were removed using Protein A Sepharose.

#### Protein expression and purification

The extracellular domain (ECD) (1-1208 a.a) was cloned into the pCAG vector (Invitrogen) with two proline substitutions at residues 986 and 987, a “GSAS” substitution at residues 682 to 685 and a C-terminal T4 fibritin trimerization motif followed by one Flag tag. The mutants were generated with a standard two-step PCR-based strategy.

The recombinant S-ECD protein (Genebank ID: QHD43416.1) was overexpressed using the HEK 293F mammalian cells (Invitrogen) at 37℃ under 5% $CO_2$ in a Multitron-Pro shaker (Infors, 130 rpm). When the cell density reached 2.0 $\times 10^6$ cells/mL, the plasmid was transiently transfected into the cells. To transfect one liter of cell culture, about 1.5 mg of the plasmid was premixed with 3 mg of polyethylenimines (PEIs) (Polysciences) in 50 mL of fresh medium for 15 mins before adding to cell culture. Cells were removed by centrifugation at 4000×g for 15 mins after sixty hours transfection. The secreted S-ECD proteins were purified by anti-FLAG M2 affinity resin (Sigma Aldrich). After loading two times, the anti-FLAG M2 resin was washed with the wash buffer containing 25 mM Tris (pH 8.0), 150 mM NaCl. The protein was eluted with the wash buffer plus 0.2 mg/mL flag peptide. The eluent was then concentrated and subjected to size-exclusion chromatography (Superose 6 Increase

10/300 GL, GE Healthcare) in buffer containing 25 mM Tris (pH 8.0), 150 mM NaCl. The peak fractions were collected and concentrated to incubate with Ab or Fab. The purified S-ECD was mixed with the Ab or Fab at a molar ratio of about 1:3.6 for one hour. Then the mixture was subjected to size-exclusion chromatography (Superose 6 Increase 10/300 GL, GE Healthcare) in buffer containing 25 mM Tris (pH 8.0), 150 mM NaCl. The peak fractions were collected for EM analysis.

**Antibody binding kinetics and competition with receptor ACE2 measured by SPR**. The binding kinetics and affinity of mAbs to SARS-CoV-2 RBD were analyzed by SPR (Biacore T200, GE Healthcare). Specifically, purified RBDs were covalently immobilized to a CM5 sensor chip via amine groups in 10 mM sodium acetate buffer (pH 5.0) for a final RU around 250. SPR assays were run at a flow rate of 30 μl/min in HEPES buffer. The sensograms were fit in a 1:1 binding model with BIA Evaluation software (GE Healthcare). To determine competition with the human ACE2 peptidase domain, SARS-CoV-2 RBD was immobilized to a CM5 sensor chip via amine group for a final RU around 250. Antibodies (1 μM) were injected onto the chip until binding steady-state was reached. ACE2 (2 μM) was then injected for 60 seconds. Blocking efficacy was determined by comparison of response units with and without prior antibody incubation.

**Neutralization of pseudotype and live virus**

SARS-CoV-2 pseudoviruses were generated by co-transfecting HEK 293T cells (ATCC) with human immunodeficiency virus backbones expressing firefly luciferase (pNL43R E luciferase) and pcDNA3.1 (Invitrogen) expression vectors encoding S proteins. Viral supernatants were collected 48h later. Pseudoviruses were incubated with serial dilutions of nAbs or Fab proteins at 37 °C for 1 h. Huh7 were then added in duplicate to the mixture. Antibody neutralization percentages were determined by measuring luciferase activity in relative light units (Bright-Glo Luciferase Assay Vector System, Promega Bioscience) 48 h after exposure to virus-antibody mixture using GraphPad Prism 7 (GraphPad Software Inc.). SARS-CoV-2 live virus focus reduction neutralization test (FRNT) was performed in a certified Biosafety level 3 laboratory as previously described. Neutralization assays against live SARS-CoV-2 were conducted using a clinical isolate (Beta/Shenzhen/SZTH-003/2020, EPI_ISL_406594 at GISAID) previously obtained from a nasopharyngeal swab of an infected patient. Serial dilutions of testing antibodies were mixed with 50 μL of SARS-CoV-2 (100 focus forming unit) in 96-well microwell plates and incubated at 37 °C for 1 h. Mixtures were then transferred to 96-well plates seeded with Vero E6 cells and allowed absorption for 1 h at 37°C. Inoculums were then removed before adding the overlay media (100 μL MEM containing 1.6% Carboxymethylcellulose). The plates were then incubated at 37°C for 24 h. Overlays were removed and then cells were fixed with 4% paraformaldehyde solution for 30 min, permeabilized with Perm/Wash buffer (BD Biosciences) containing 0.1% Triton X-100 for 10 min. Cells were incubated with rabbit anti-SARS-CoV-2 NP IgG (Sino Biological, Inc) for 1 h at room temperature before adding HRP-conjugated goat anti-rabbit IgG (H+L) antibody (TransGen Biotech, Beijing). The reactions were developed with KPL TrueBlue Peroxidase substrates (Seracare Life Sciences Inc). The numbers of SARS-CoV-2 foci were calculated using an EliSpot reader (Cellular Technology Ltd).

**Shedding of S1 from cell surface expressed SARS-CoV-2 S glycoprotein**

Plasmids encoding SARS-CoV-2 S or mutant S containing GSAS, substituting RRAR at the junction between S1 and S2 to avoid digestion by Furin protease protein on the cell surface were transfected into HEK293T cells. Cells Samples were prepared in multiples for serial incubations at 37 °C for 120, 60, 45, 30, 15, or 5 min. Immediately after the allocated incubation time, antibody-stained cells were transferred to ice then thoroughly washed with ice-cold PBS and 2% FBS. Samples were then stained with anti-human IgG Fc PE (Biolegend 410718) for nAbs, or anti-human IgG (H+L) Alexa Flour 647 (ThermoFisher A21445) for Fab. After thorough washes with ice-cold PBS and 2% FBS, samples were resuspended and analyzed with FACS Calibur (BD Biosciences, USA) and FlowJo 10 software (FlowJo, USA). Binding at each of the allocated time points was determined by the MFI weighted by multiplying the number of positive cells in the selected gates and normalized in relative to that at the 5 min time point.

**Cryo-EM sample preparation**

The peak fractions of complex was concentrated to about 1.5 mg/mL and applied to the grids. Aliquots (3.3 μL) of the protein complex were placed on glow-discharged holey carbon grids (Quantifoil Au R1.2/1.3). The grids were blotted for 2.5 s or 3.0 s and flash-frozen in liquid ethane cooled by liquid nitrogen with Vitrobot (Mark IV, Thermo Scientific). The cryo-EM samples were transferred to a Titan Krios operating at 300 kV equipped with Cs corrector, Gatan K3 Summit detector and GIF Quantum energy filter. Movie stacks were automatically collected using AutoEMation [1], with a slit width of 20 eV on the energy filter and a defocus range from -1.2 µm to -2.2 µm in super-resolution mode at a nominal magnification of 81,000×. Each stack was exposed for 2.56 s with an exposure time of 0.08 s per frame, resulting in a total of 32 frames per stack. The total dose rate was approximately 50 $e^-/Å^2$ for each stack. The stacks were motion corrected with MotionCor2 [2] and binned 2-fold, resulting in a pixel size of 1.087 Å/pixel. Meanwhile, dose weighting was performed [3]. The defocus values were estimated with Gctf [4].

**Data processing**

Particles for all samples were automatically picked using Relion 3.0.6 [5-8] from manually selected micrographs. After 2D classification with Relion, good particles were selected and subject to two cycle of heterogeneous refinement without symmetry using cryoSPARC [9].The good particles were selected and subjected to Non-uniform Refinement (beta) with C1 symmetry, resulting in the 3D reconstruction for the whole structures, which was further subject to 3D auto-refinement and post-processing with Relion. For interface between SARS-CoV-2 S protein and all kinds of mAb, the dataset was subject to focused refinement with adapted mask on each RBD-mAb sub-complex to improve the map quality. Then the dataset of three similar RBD-mAb sub-complexes were combined and subject to focused refinement with Relion. And the combined dataset was re-centered on the interface between RBD and mAb and re-extracted. The re-extracted dataset was 3D classified with Relion focused on RBD-mAb sub-complex. Then the good particles were selected and subject to focused refinement with Relion, resulting in the 3D reconstruction of better quality on RBD-mAb sub-complex.

The resolution was estimated with the gold-standard Fourier shell correlation 0.143 criterion [10] with high-resolution noise substitution [11]. Refer to Supplemental Figures S4-S5 and Supplemental Table S3 for details of data collection and processing.

**Model building and structure refinement**

For model building of all complex of S-ECD of SARS-CoV-2 with all mAb, the atomic model (PDB ID: 7C2L) were used as templates, which were molecular dynamics flexible fitted [12] into the whole cryo-EM map of the complex and the focused-refined cryo-EM map of the RBD-mAb sub-complex, respectively. And the fitted atomic models were further manually adjusted with Coot [13]. Each residue was manually checked with the chemical properties taken into consideration during model building. Several segments, whose corresponding densities were invisible, were not modeled. Structural refinement was performed in Phenix [14] with secondary structure and geometry restraints to prevent overfitting. To monitor the potential overfitting, the model was refined against one of the two independent half maps from the gold-standard 3D refinement approach. Then, the refined model was tested against the other map. Statistics associated with data collection, 3D reconstruction and model building were summarized in Table S3.

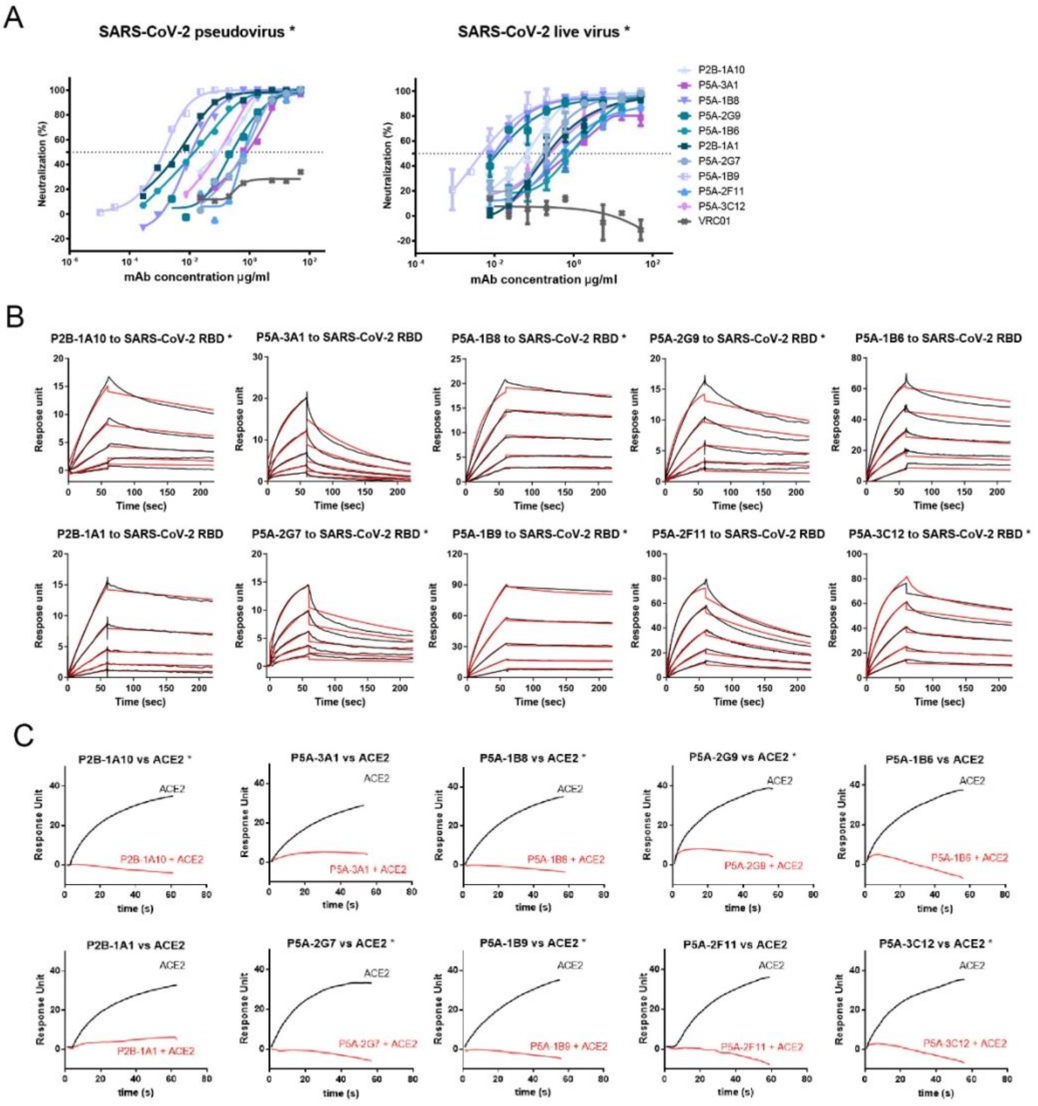


**Fig. S1 | Neutralizing activity, binding kinetics, and ACE2 competition of COVID-19 donor-derived neutralizing mAbs.**

**(A)**, Neutralizing activities of the antibodies against SARS-Cov-2 pseudovirus and live virus. (**B)**, binding kinetics of antibodies with soluble SARS-Cov-2 RBD measured by Surface Plasmon Resonance. The black lines indicate the experimentally derived curves while the red lines represent fitted curves by 1:1 binding fitness model. (**C)**, binding patterns of ACE2 receptor protein to SARS-Cov-2 RBD with (red curve) or without (black curve) prior injection and saturation with each testing antibody.

* Published in the reference (Zhang, et al. Public neutralizing antibodies elicited by SARS-CoV-2 infection. submitted).

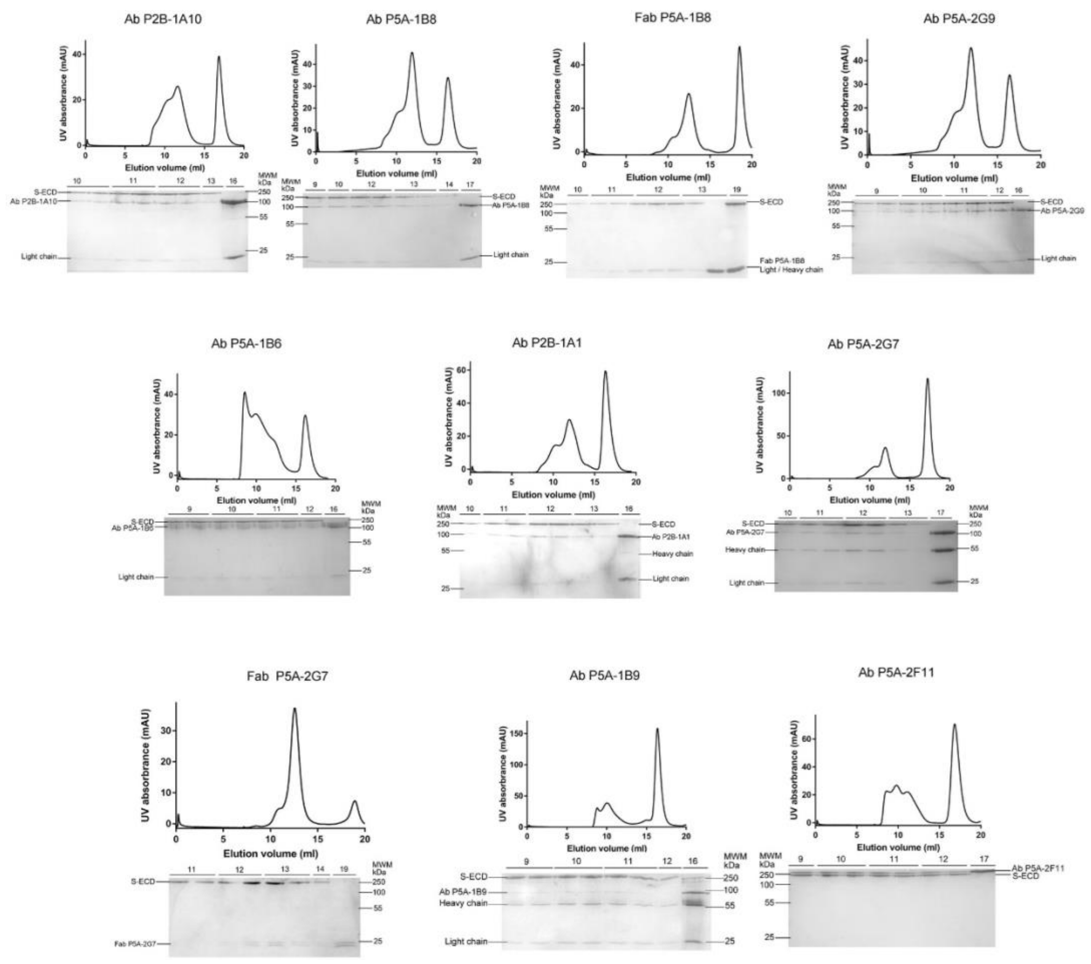


**Fig. S2 | Representative SEC purification profile of the S-ECD of SARS-CoV-2 in complex with all kinds of mAb.**

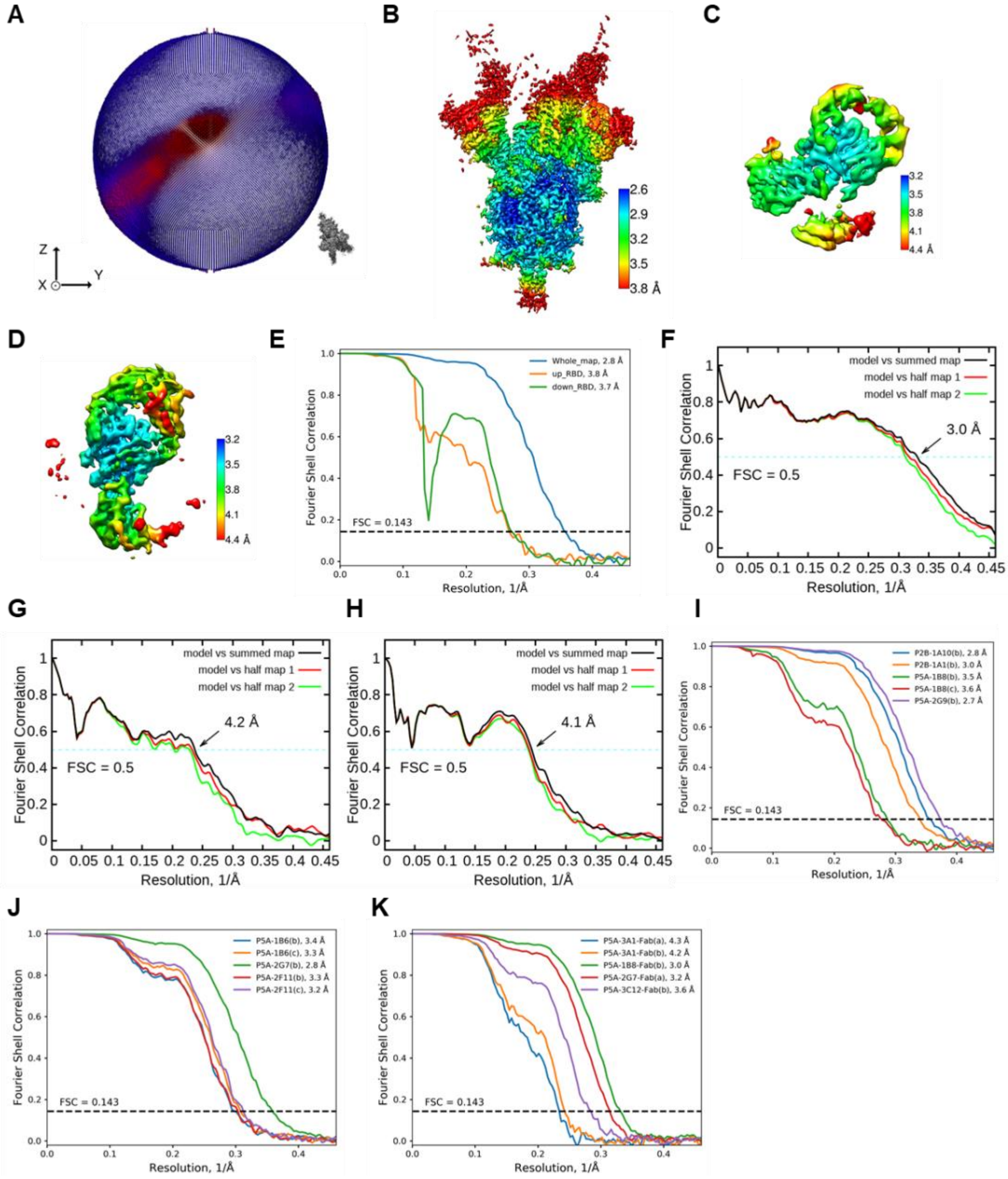


**Fig. S3 | Representative Cryo-EM analysis of S-ECD in complex with P5A-1B9.** **(A),** Euler angle distribution in the final 3D reconstruction of S-ECD of SARS-CoV-2 bound with P5A-1B9 complex. **(B)-(D),** Local resolution map for the 3D reconstruction of overall structure, "down" RBD-P5A-1B9 sub-complex and "up" RBD-P5A-1B9 sub-complex, respectively. **(E),** FSC curve of the overall structure (blue), "up" RBD-P5A-1B9 sub-complex (orange) and "down" RBD-P5A-1B9 sub-complex (green). **(F),** FSC curve of the refined model of SARS-CoV-2 bound with P5A-1B9 complex versus the overall structure that it is refined against (black); of the model refined against the first half map versus the same map (red); and of the model refined against the first half map versus the second half map (green). The small difference between the red and green curves indicates that the refinement of the atomic coordinates did not suffer from overfitting. (**G)** and **(H),** FSC curve of the refined model of “down” RBD-P5A-1B9 sub-complex and “up” RBD-P5A-1B9 sub-complex, which is same to the (**H)**. (**I)-(K),** Gold standard FSC curve of the overall structure for all kinds of mAb bound with S-ECD, respectively. (a), mono; (b) double;(c), triple.

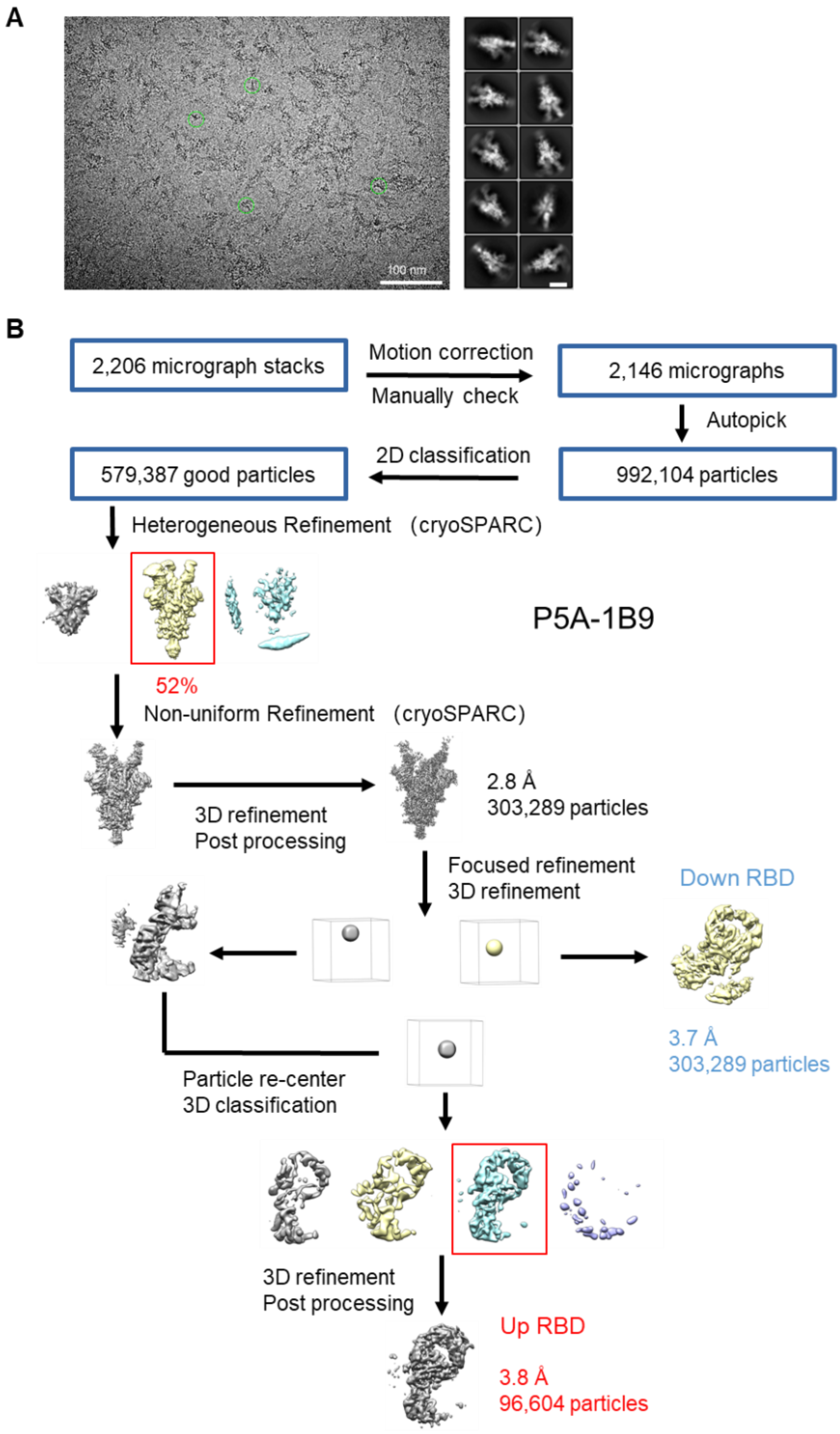


**Fig. S4 | Representative cryo-EM image and flowchart for cryo-EM data processing of S-ECD in complex with P5A-1B9.**

**(A),** Representative electron micrograph and 2D class averages of cryo-EM particle images. The scale bar in 2D class averages represent 10 nm. **(B),** Please refer to the 'Data Processing' in Methods section for details.

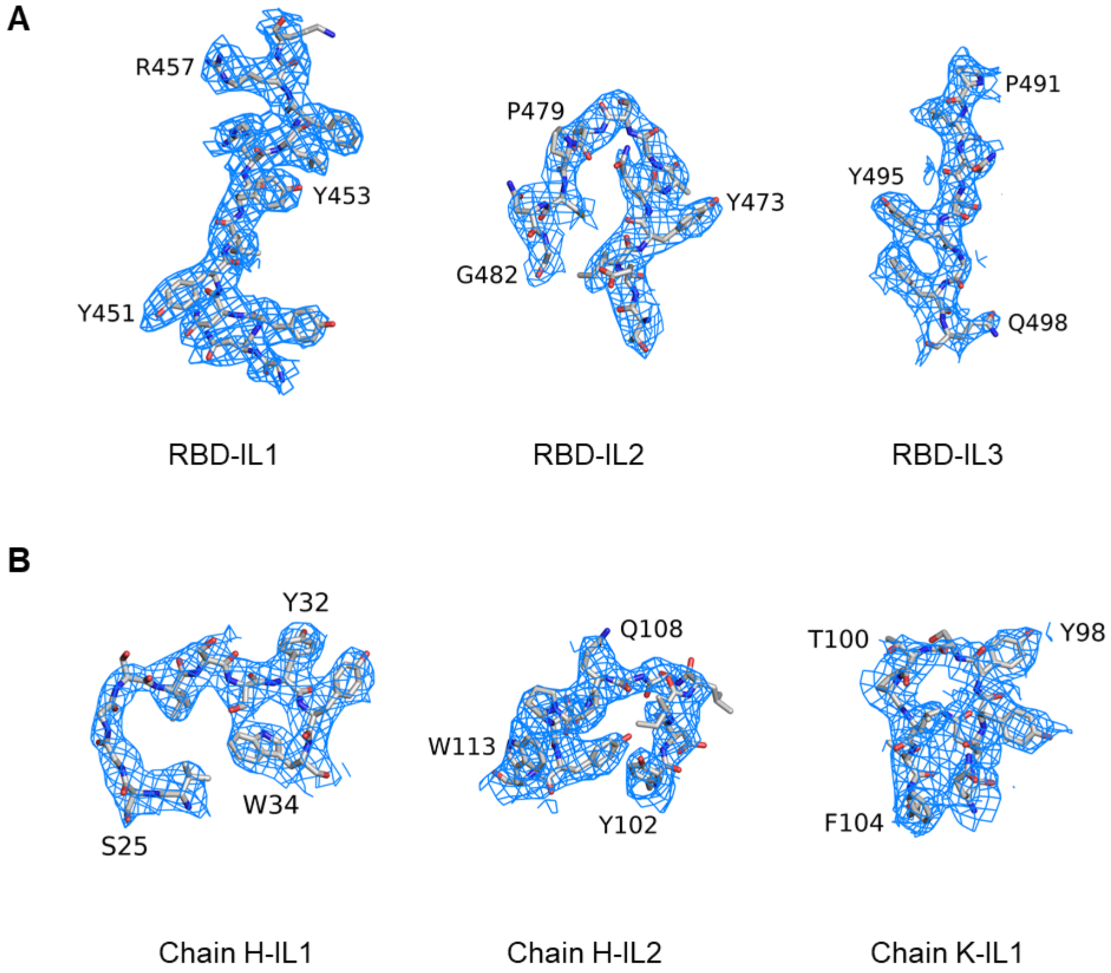


**Fig. S5 | Representative cryo-EM density maps. (A),** Cryo-EM density map for RBD of S-ECD in complex with P5A-1B9 shown at threshold of 8 σ. **(B),** Cryo-EM density map for mAb of S-ECD in complex with P5A-1B9 shown at threshold of 8 σ. IL, interface loop.

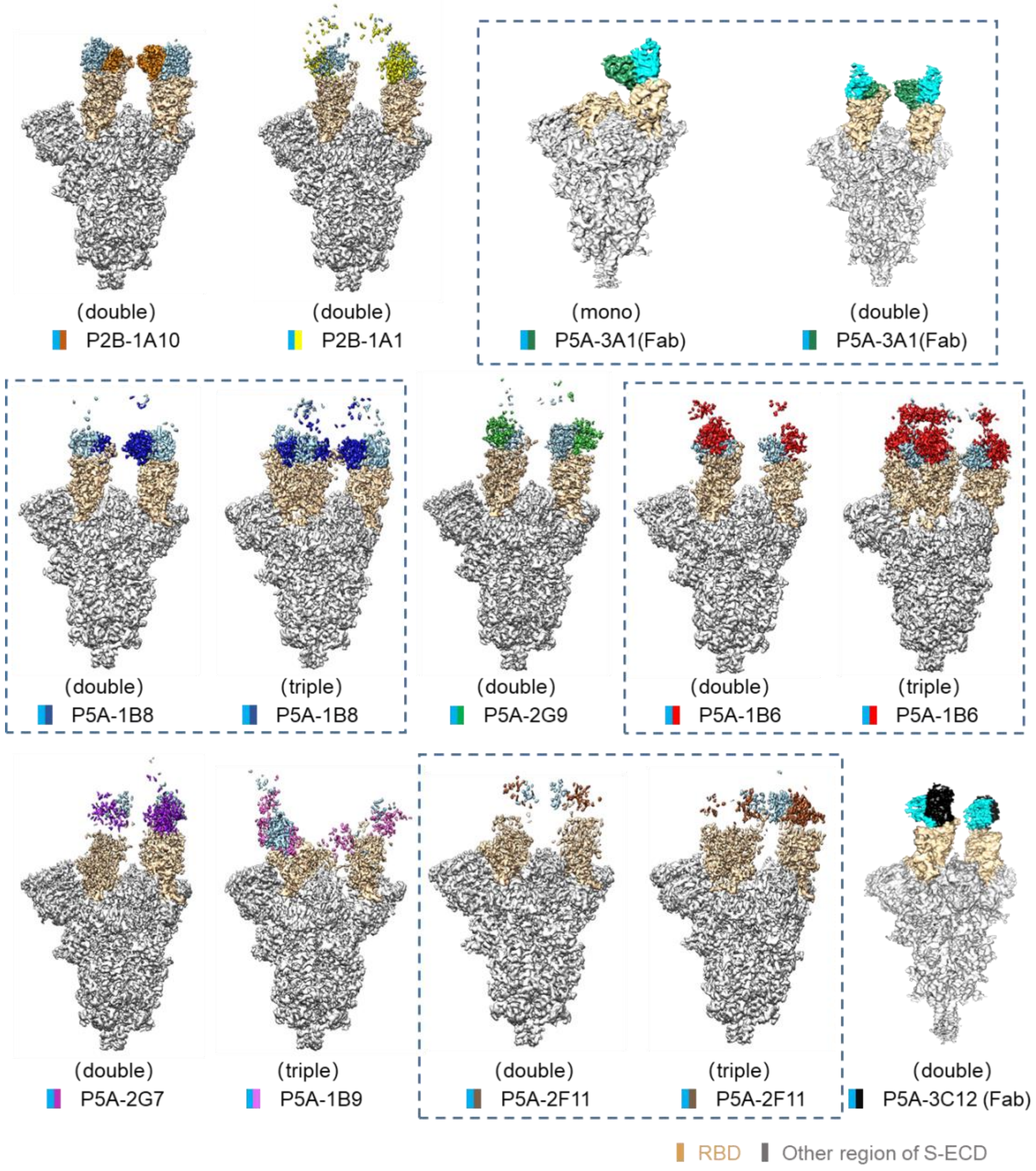


**Fig. S6 | All cryo-EM maps of S-ECD in complex with mAb.**

The domain-colored cryo-EM maps of the all complex are shown here.

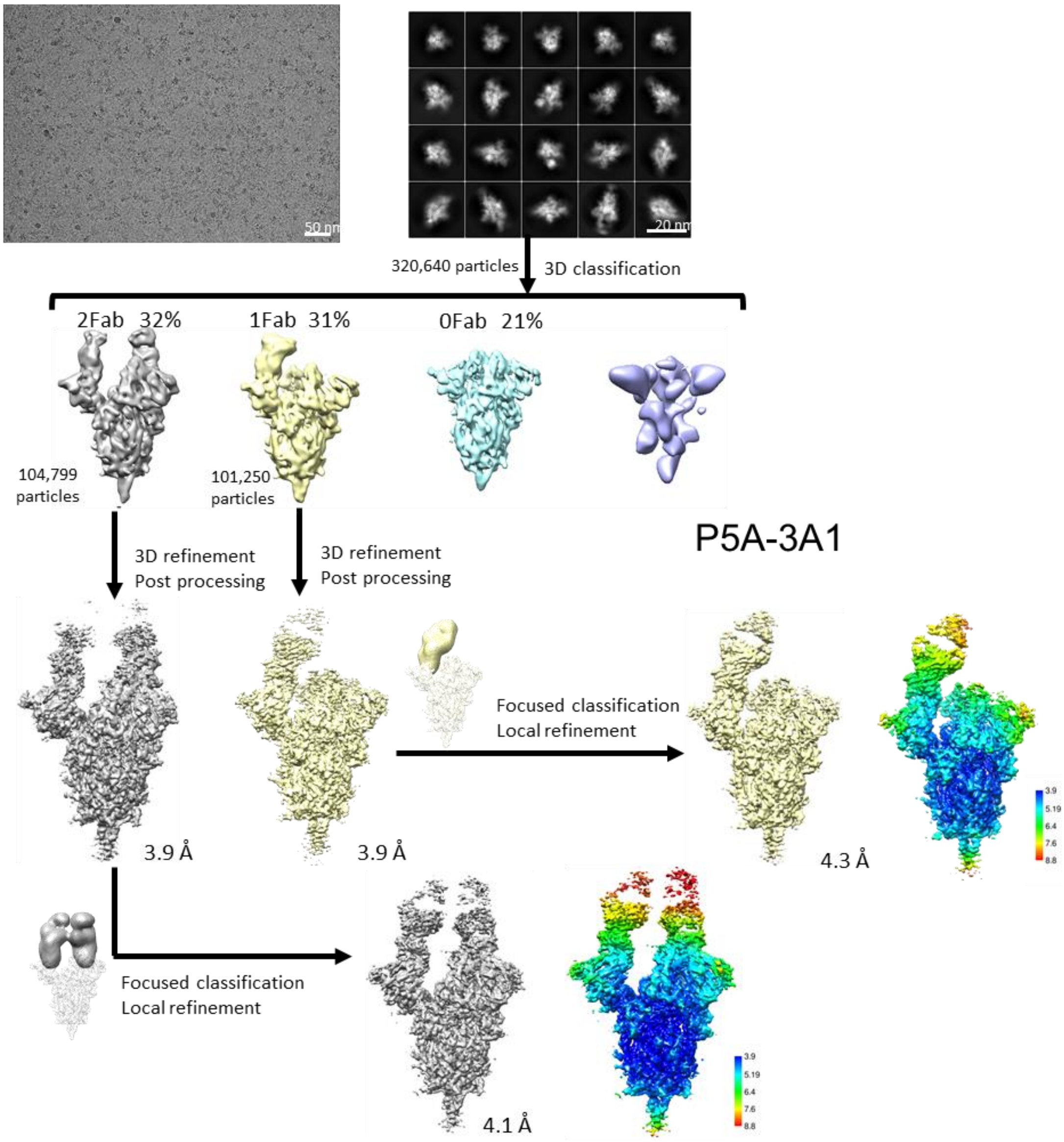


**Fig. S7 | Flowchart for cryo-EM data processing of S-ECD in complex with P5A-3A1.**

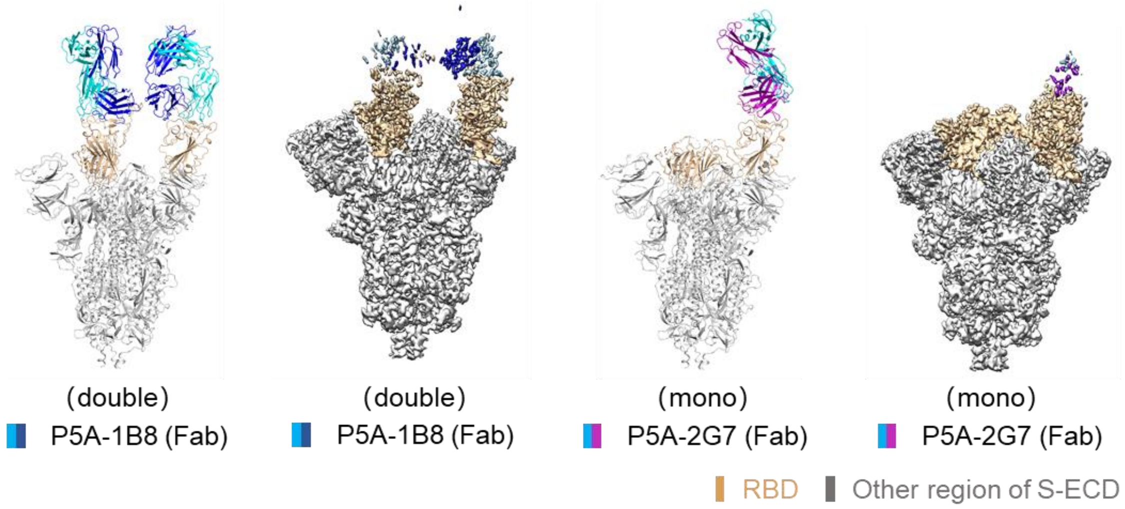


**Fig. S8 | All cryo-EM models and maps of S-ECD in complex with Fab.**

The domain-colored models and cryo-EM maps of all complex are shown here.

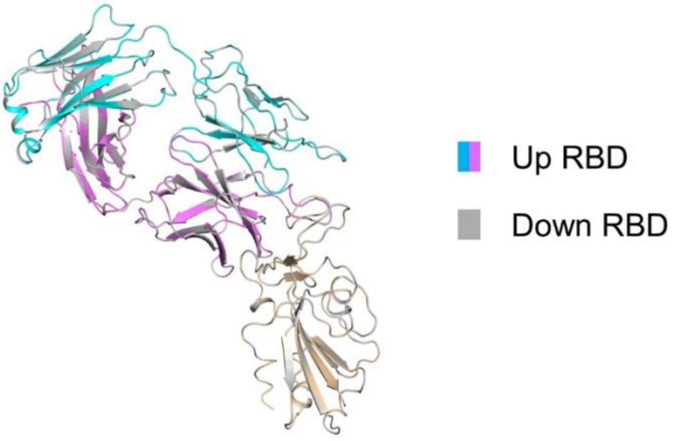


**Fig. S9 | Structural alignment of RBD-P5A-1B9 sub-complex.**

Superposition in local structure of “up” RBD-P5A-1B9 sub-complex and “down” RBD-P5A-1B9 sub-complex, which has no difference between two maps.

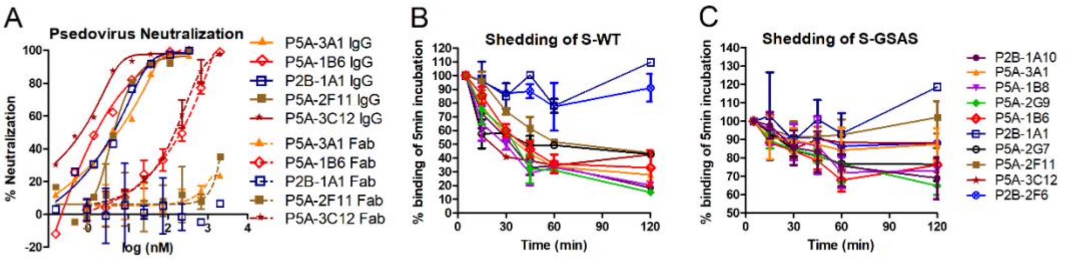


**Fig. S10 | The neutralization and shedding property of rest antibodies.**

**(A)**, Neutralizing activity against SARS-CoV-2 psedovirus by rest IgG forms (solid line) and Fab forms (dotted line). Shedding of S1 over time was measured by flow cytometry at 37°C with 293T cell-surface expressed wildtype SARS-Cov-2 spike (B) or a mutant spike containing GSAS substitution at S1/S2 cleavage motif (C). The percentage of cells at each allocated time point was determined by the MFI weighted by multiplying the number of positive cells in the selected gates and normalized in relative to the 5 min time point. Data shown were from at least two independent experiments. Values are indicated as mean±SEM.

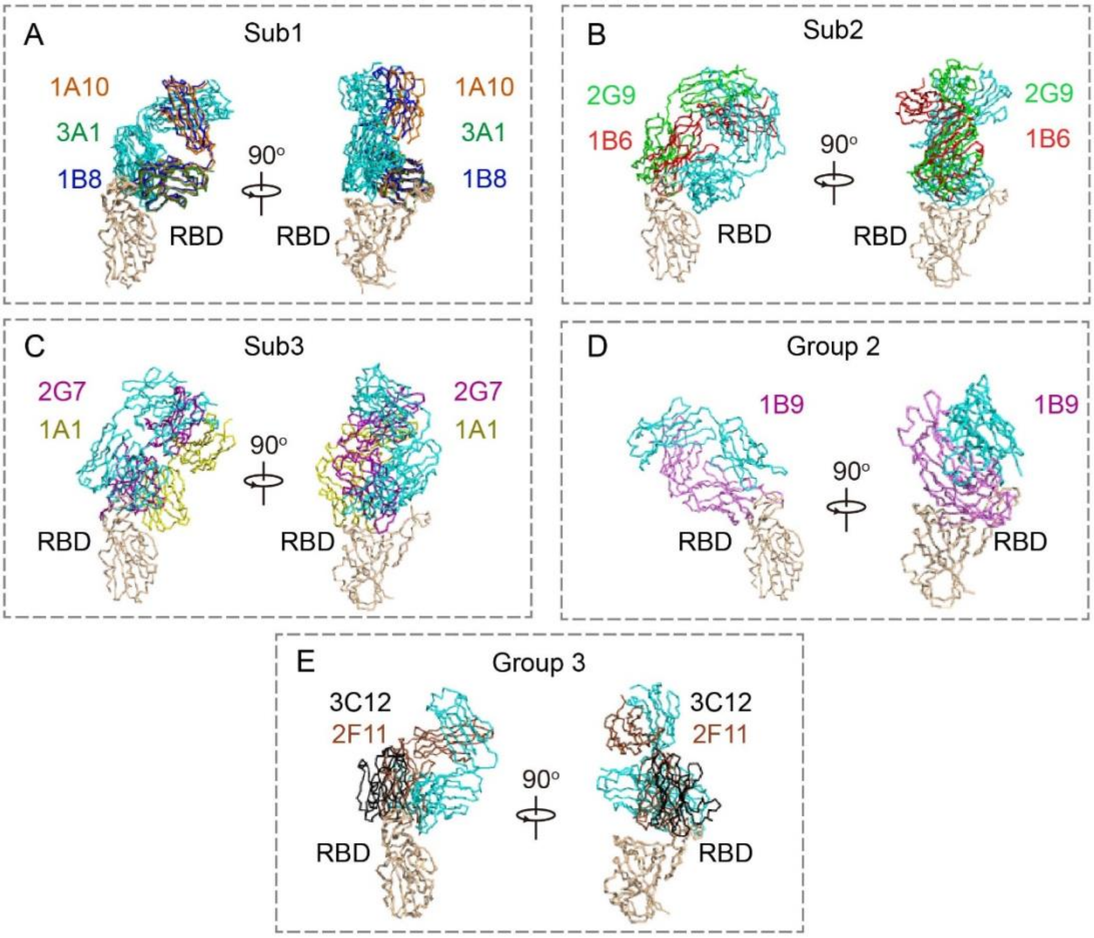


**Fig. S11 | Classification and exhibition of the 10 antibodies.** The 10 antibodies could be classified into three groups. The first group antibodies can be further divided into three subgroups. All RBD-Ab complexes are shown as ribbon mode in two vertical directions. The color mode of our 10 antibodies are the same as Fig 4. **(A)** Antibodies of subgroup 1. Subgroup 1 consists of 3 antibodies P2B-1A10, P5A-3A1 and P5A-1B8. **(B)** Antibodies of subgroup 2. Subgroup 2 consists of 2 antibodies P5A-2G9 and P5A-1B6. **(C)** Antibodies of subgroup 3. Subgroup 3 consists of 2 antibodies P2B-1A1 and P5A-2G7. **(D)** Antibody of group 2. Group 2 consist of 1 antibody P5A-1B9. **(E)** Antibodies of group 3. Group 3 consists of 2 antibodies P5A-2F11 and P5A-3C12.

**Supplemental Table 1 | Gene family analysis of COVID-19 donor-derived neutralizing mAbs**

| | Heavy chain | | | | Kappa chain (K) or Lambda chain (L) | | | |
|---|---|---|---|---|---|---|---|---|
| mAbs | IGHV | HCDR3 | HCDR3 length | SHM (%) | IGK(L)V | K(L)CDR3 | K(L)CDR3 length | SHM (%) |
| P2B-1A1 | 4-59*01 | ARLERDWPLDAFDI | 14 | 0.35 | L2-14*01 | SSYTSNNTFA | 10 | 1.11 |
| P2B-1A10* | 3-53*01 | AREGPKSITGTAFDI | 15 | 0.35 | K1-33*01,K1D-33*01 | QQYDNLPMYT | 10 | 0.38 |
| P5A-3A1 | 3-53*01 | ARDYGDFYFDY | 11 | 0.00 | K3-20*01 | QQYGSSPRT | 9 | 0.00 |
| P5A-1B8* | 3-53*01 | ARETLAFDY | 9 | 1.40 | K1-9*01 | QQLNSYPPA | 9 | 0.00 |
| P5A-2G9* | 3-33*01,3-33*06 | ARWFHTGGYFDY | 12 | 0.00 | L5-37*01 | MIWPSNALYV | 10 | 0.35 |
| P5A-1B6 | 3-30*04,3-30-3*03 | ARDGQAITMVQGVIGPPFDY | 20 | 0.00 | K1-33*01,K1D-33*01 | QQYDNLPYT | 9 | 0.00 |
| P5A-2G7* | 4-61*01 | ARERCYYGSGRAPRCVWFDP | 20 | 0.34 | L2-14*01 | SSYTSSSTLVV | 11 | 0.74 |
| P5A-1B9* | 4-59*01 | ASNGQYYDILTGQPPDYWYFD | 22 | 0.70 | K4-1*01 | QQYYSTPLT | 9 | 0.00 |
| P5A-2F11 | 1-8*01 | ARYIVVVPAAKGFDP | 15 | 0.00 | K4-1*01 | QQYYSTPLT | 9 | 0.00 |
| P5A-3C12* | 2-5*02 | AHSLFLTVGYSSSWSPFDY | 19 | 0.00 | K4-1*01 | QQYYSTPHT | 9 | 0.00 |

The program IMGT/V-QUEST was applied to analyze gene germline, complementarity determining region (CDR) 3 length, and somatic hyper mutation (SHM). The SHM frequency was calculated from the mutated nucleotides.

* Published in the reference (Zhang, et al. Public neutralizing antibodies elicited by SARS-CoV-2 infection. submitted)

### Supplemental Table 2 | Data collection, 3D reconstruction and model statistic

| | | | | | | |
|---|---|---|---|---|---|---|
| **Data collection** | | | | | | |
| EM equipment | Titan Krios (Thermo Fisher Scientific) | | | | | |
| Voltage (kV) | 300 | | | | | |
| Detector | Gatan K3 Summit | | | | | |
| Energy filter | Gatan GIF Quantum, 20 eV slit | | | | | |
| Pixel size (Å) | 1.087 | | | | | |
| Electron dose (e-/Å2) | 50 | | | | | |
| Defocus range (μm) | -1.2 ~ -2.2 | | | | | |
| Number of collected micrographs | 2,522 | 605 | | 2,477 | 852 | |
| Number of selected micrographs | 2,440 | 566 | | 2,356 | 796 | |
| Sample | P2B-1A10 | P5A-1B8[b] | P5A-1B8[c] | P2A-2G9 | P5A-1B6[b] | P5A-1B6[c] |
| PDB code | 7CZQ | 7CZR | 7CZS | 7CZT | 7CZU | 7CZV |
| EMDB code | EMD-30513 | EMD-30514 | EMD-30515 | EMD-30516 | EMD-30517 | EMD-30518 |
| **3D Reconstruction** | | | | | | |
| Software | cryoSPARC/ Relion | | | | | |
| Number of used particles | 430,062 | 26,128 | 16,489 | 497,874 | 39,454 | 55,619 |
| Resolution (Å) | 2.8 | 3.5 | 3.6 | 2.7 | 3.4 | 3.3 |
| Symmetry | C1 | | | | | |
| Map sharpening B factor ($Å^2$) | -90 | | | | | |
| **Refinement** | | | | | | |
| Software | Phenix | | | | | |
| Cell dimensions (Å) | 313.056 | | | | | |
| Model composition | | | | | | |
| Protein residues | 3,868 | 3,854 | 4310 | 3872 | 3878 | 4346 |
| Side chains assigned | 3,868 | 3,854 | 4310 | 3872 | 3878 | 4346 |
| Sugar | 70 | 70 | 71 | 70 | 70 | 71 |
| R.m.s deviations | | | | | | |
| Bonds length (Å) | 0.008 | 0.006 | 0.006 | 0.006 | 0.006 | 0.006 |
| Bonds Angle (˚) | 0.933 | 0.876 | 0.877 | 0.888 | 0.814 | 1.858 |
| Ramachandran plot statistics (%) | | | | | | |
| Favored | 92.74 | 93.11 | 90.99 | 93.79 | 93.52 | 92.95 |
| Allowed | 7.23 | 6.86 | 8.91 | 6.17 | 6.45 | 6.92 |
| Outlier | 0.03 | 0.03 | 0.10 | 0.03 | 0.03 | 0.13 |

[a], mono; [b], double; [c], triple

### Supplemental Table 2, continued

| | | | | | |
|---|---|---|---|---|---|
| **Data collection** | | | | | |
| EM equipment | Titan Krios (Thermo Fisher Scientific) | | | | |
| Voltage (kV) | 300 | | | | |
| Detector | Gatan K3 Summit | | | | |
| Energy filter | Gatan GIF Quantum, 20 eV slit | | | | |
| Pixel size (Å) | 1.087 | | | | |
| Electron dose (e-/Å2) | 50 | | | | |
| Defocus range (μm) | -1.2 ~ -2.2 | | | | |
| Number of collected micrographs | 892 | 1,343 | 2,206 | 920 | |
| Number of selected micrographs | 863 | 1,282 | 2,146 | 897 | |
| Sample | P2B-1A1 | P5A-2G7 | P5A-1B9 | P5A-2F11[b] | P5A-2F11[c] |
| PDB code | 7CZP | 7CZW | 7CZX | 7CZY | 7CZZ |
| EMDB code | EMD-30512 | EMD-30519 | EMD-30520 | EMD-30521 | EMD-30522 |
| **3D Reconstruction** | | | | | |
| Software | cryoSPARC/ Relion | | | | |
| Number of used particles | 146,875 | 211,771 | 303,289 | 39,337 | 55,704 |
| Resolution (Å) | 3.0 | 2.8 | 2.8 | 3.3 | 3.2 |
| Symmetry | C1 | | | | |
| Map sharpening B factor ($Å^2$) | -90 | | | | |
| **Refinement** | | | | | |
| Software | Phenix | | | | |
| Cell dimensions (Å) | 313.056 | | | | |
| Model composition | | | | | |
| Protein residues | 3862 | 3880 | 4,368 | 3,880 | 4349 |
| Side chains assigned | 3862 | 3880 | 4,368 | 3,880 | 4349 |
| Sugar | 74 | 70 | 71 | 76 | 80 |
| R.m.s deviations | | | | | |
| Bonds length (Å) | 0.009 | 0.009 | 0.005 | 0.008 | 0.007 |
| Bonds Angle (˚) | 0.994 | 1.01 | 0.897 | 0.907 | 0.893 |
| Ramachandran plot statistics (%) | | | | | |
| Favored | 92.26 | 91.78 | 92.44 | 92.74 | 92.14 |
| Allowed | 7.67 | 8.19 | 7.39 | 7.23 | 7.69 |
| Outlier | 0.07 | 0.03 | 0.17 | 0.03 | 0.17 |

[a], mono; [b], double; [c], triple

### Supplemental Table 2, continued

| | | | | | |
|---|---|---|---|---|---|
| **Data collection** | | | | | |
| EM equipment | Titan Krios (Thermo Fisher Scientific) | | | | |
| Voltage (kV) | 300 | | | | |
| Detector | Gatan K3 Summit | | | | |
| Energy filter | Gatan GIF Quantum, 20 eV slit | | | | |
| Pixel size (Å) | 1.087 | | | | |
| Electron dose (e-/Å2) | 50 | | | | |
| Defocus range (μm) | -1.2 ~ -2.2 | | | | |
| Number of collected micrographs | 2,352 | | 1,215 | 1,480 | 5,155 |
| Number of selected micrographs | 2,353 | | 911 | 1,401 | 5,155 |
| Sample | P5A-3A1-Fab[a] | P5A-3A1-Fab[b] | P5A-1B8-Fab | P5A-2G7-Fab | P5A-3C12-Fab |
| PDB code | 7D0B | 7D0C | 7D00 | 7D03 | 7D0D |
| EMDB code | EMD-30529 | EMD-30530 | EMD-30523 | EMD-30524 | EMD-30531 |
| **3D Reconstruction** | | | | | |
| Software | cryoSPARC/ Relion | | | | |
| Number of used particles | 41,778 | 43,126 | 145,010 | 239,537 | 120,531 |
| Resolution (Å) | 4.3 | 4.2 | 3.0 | 3.2 | 3.6 |
| Symmetry | C1 | | | | |
| Map sharpening B factor ($Å^2$) | -90 | | | | |
| **Refinement** | | | | | |
| Software | Phenix | | | | |
| Cell dimensions (Å) | 313.056 | | | | |
| Model composition | | | | | |
| Protein residues | 3172 | 3415 | 3854 | 3412 | 2973 |
| Side chains assigned | 3172 | 3415 | 3854 | 3412 | 2973 |
| Sugar | 62 | 64 | 70 | 69 | 63 |
| R.m.s deviations | | | | | |
| Bonds length (Å) | 0.008 | 0.005 | 0.009 | 0.009 | 0.006 |
| Bonds Angle (˚) | 1.006 | 0.864 | 0.942 | 0.959 | 0.867 |
| Ramachandran plot statistics (%) | | | | | |
| Favored | 90.67 | 93.07 | 92.09 | 91.66 | 93.17 |
| Allowed | 9.33 | 6.93 | 7.84 | 8.24 | 6.8 |
| Outlier | 0.00 | 0.00 | 0.07 | 0.10 | 0.03 |

[a], mono; [b], double; [c], triple

1 Lei, J. & Frank, J. Automated acquisition of cryo-electron micrographs for single particle reconstruction on an FEI Tecnai electron microscope. *Journal of structural biology* **150**, 69-80, doi:10.1016/j.jsb.2005.01.002 (2005).

2 Zheng, S. Q. *et al.* MotionCor2: anisotropic correction of beam-induced motion for improved cryo-electron microscopy. *Nature methods* **14**, 331-332, doi:10.1038/nmeth.4193 (2017).

3 Grant, T. & Grigorieff, N. Measuring the optimal exposure for single particle cryo-EM using a 2.6 A reconstruction of rotavirus VP6. *eLife* **4**, e06980, doi:10.7554/eLife.06980 (2015).

4 Zhang, K. Gctf: Real-time CTF determination and correction. *Journal of structural biology* **193**, 1-12, doi:10.1016/j.jsb.2015.11.003 (2016).

5 Zivanov, J. *et al.* New tools for automated high-resolution cryo-EM structure determination in RELION-3. *eLife* **7**, doi:10.7554/eLife.42166 (2018).

6 Kimanius, D., Forsberg, B. O., Scheres, S. H. & Lindahl, E. Accelerated cryo-EM structure determination with parallelisation using GPUs in RELION-2. *eLife* **5**, doi:10.7554/eLife.18722 (2016).

7 Scheres, S. H. RELION: implementation of a Bayesian approach to cryo-EM structure determination. *Journal of structural biology* **180**, 519-530, doi:10.1016/j.jsb.2012.09.006 (2012).

8 Scheres, S. H. A Bayesian view on cryo-EM structure determination. *Journal of molecular biology* **415**, 406-418, doi:10.1016/j.jmb.2011.11.010 (2012).

9 Punjani, A., Rubinstein, J. L., Fleet, D. J. & Brubaker, M. A. cryoSPARC: algorithms for rapid unsupervised cryo-EM structure determination. *Nature methods* **14**, 290-296, doi:10.1038/nmeth.4169 (2017).

10 Rosenthal, P. B. & Henderson, R. Optimal determination of particle orientation, absolute hand, and contrast loss in single-particle electron cryomicroscopy. *Journal of molecular biology* **333**, 721-745 (2003).

11 Chen, S. *et al.* High-resolution noise substitution to measure overfitting and validate resolution in 3D structure determination by single particle electron

cryomicroscopy. *Ultramicroscopy* **135**, 24-35, doi:10.1016/j.ultramic.2013.06.004 (2013).

12 Trabuco, L. G., Villa, E., Mitra, K., Frank, J. & Schulten, K. Flexible fitting of atomic structures into electron microscopy maps using molecular dynamics. *Structure (London, England : 1993)* **16**, 673-683, doi:10.1016/j.str.2008.03.005 (2008).

13 Emsley, P., Lohkamp, B., Scott, W. G. & Cowtan, K. Features and development of Coot. *Acta crystallographica. Section D, Biological crystallography* **66**, 486-501, doi:10.1107/S0907444910007493 (2010).

14 Adams, P. D. *et al.* PHENIX: a comprehensive Python-based system for macromolecular structure solution. *Acta crystallographica. Section D, Biological crystallography* **66**, 213-221, doi:10.1107/S0907444909052925 (2010).
